# Synergy of calcium release site determinants in control of calcium release events in cardiac myocytes

**DOI:** 10.1101/2020.08.26.260968

**Authors:** B. I. Iaparov, I. Zahradnik, A. S. Moskvin, A. Zahradnikova

## Abstract

Recent data on structure of dyads in cardiac myocytes indicate variable clustering of RyR calcium release channels. The question arises as to how geometric factors of RyR arrangement translate to their role in formation of calcium release events (CRE). Since this question is not experimentally testable *in situ*, we performed *in silico* experiments on a large set of calcium release site (CRS) models. The models covered the range of RyR spatial distributions observed in dyads, and included gating of RyRs with open probability dependent on Ca^2+^ and Mg^2+^ concentration. The RyR single-channel calcium current, varied in the range of previously reported values, was set constant in the course of CRE simulations. Other known features of dyads were omitted in the model formulation for clarity. CRE simulations initiated by a single random opening of one of the RyRs in a CRS produced spark-like responses with characteristics that varied with RyR vicinity, a newly defined parameter quantifying spatial distribution of RyRs in the CRSs, and with the RyR single-channel calcium current. The CRE characteristics followed the law of mass action with respect to a CRS state variable, defined as a weighed product of RyR vicinity and RyR single-channel calcium current. The results explained the structure-function relations among determinants of cardiac dyads on synergy principles and thus allowed to evolve the concept of CRS as a dynamic unit of cardiac dyad.

## Introduction

Calcium sparks are miniature transient increases of fluorescence intensity observable in the cytosol of cardiac myocytes by means of calcium-sensitive fluorescent dyes (Cheng et al., 1993; Lopez-Lopez et al., 1994). They represent transient elementary calcium release events (CREs) appearing locally at dyads, the intracellular structural complexes of cardiac myocyte, where terminal cisternae of sarcoplasmic reticulum connect to sarcolemma (Sun et al., 1995; Franzini-Armstrong et al., 1998). During excitation of a cardiac myocyte these events are triggered in synchrony over the whole volume of a myocyte by influx of Ca^2+^ ions via sarcolemmal Cav1.2 calcium channels (Cannell et al., 1995; Lopez-Lopez et al., 1995), and thus provide myofibrillar sarcomeres with calcium ions needed for contraction (Eisner et al., 2017).

Elementary CREs may also occur spontaneously in myocytes of resting myocardium. Under physiological conditions these spontaneous CREs occur at low frequency; however, under pathological conditions their frequency may increase and thus initiate arrhythmogenic calcium waves (Cheng et al., 1996; Lukyanenko and Gyorke, 1999). Spontaneous CREs originate at the terminal cisternae of sarcoplasmic reticulum, where ryanodine receptor calcium-release channels (RyRs) form large clusters (Franzini-Armstrong et al., 1999). Local CREs depend on RyR activity, which is regulated by cytosolic and luminal Ca^2+^ ions, cytosolic Mg^2+^, cytosolic ATP^2-^, and by numerous regulatory proteins and processes.

The structure of dyads studied by electron microscopy revealed common features in their construction (Franzini-Armstrong et al., 1999; Hayashi et al., 2009) as well as substantial structural variability (Novotova et al., 2009; Novotova et al., 2013; Novotova et al., 2020). Super-resolution localization studies revealed tendency of RyRs to form irregular clusters of variable size and surface density (Baddeley et al., 2009; Hou et al., 2015; Macquaide et al., 2015; Galice et al., 2018; Jayasinghe et al., 2018; Kolstad et al., 2018; Asghari et al., 2020). Moreover, the composition of CRSs may change in parallel with changes in the structure of dyads (Asghari et al., 2020; Drum et al., 2020; Novotova et al., 2020). Thus, the understanding of the dyad as the source of local CREs became ambiguous for the difficulty to define explicitly the common source structure. Rather, the dyads can be seen as the morphological substrate of calcium release sites, where calcium release events emerge. The function of dyad as a calcium release site (CRS) can be probed only indirectly. It is performed by monitoring the local calcium release events as calcium sparks or as calcium spikes (Cheng et al., 1993; Song et al., 1998). Despite the recent progress in technologies, these approaches have substantial technical limitations mainly due to the low spatio-temporal resolution and low signal-to-noise ratio, inherent to the calcium-sensitive fluorescent dyes (Zahradnikova et al., 2007; Kong et al., 2008; Janicek et al., 2013). Interpretation of local fluorescent signals is also problematic, since it is based on data measured indirectly or in artificial systems. The amplitude and kinetics of CREs varies broadly and includes not only sparks but also the invisible quarks (Lipp and Niggli, 1996), “quarky calcium release” (Brochet et al., 2011) and prolonged sparks (Zima et al., 2008; Brochet et al., 2011), and may vary even if recorded at the same spot (Sobie et al., 2006; Zima et al., 2008; Janicek et al., 2012). Therefore, theoretical approaches could be of help to formulate and verify working hypotheses (Soeller and Cannell, 2004; Williams et al., 2007; Soeller et al., 2009; Williams et al., 2011; Stern et al., 2013).

According to recent understanding, calcium sparks result from group activity of numerous RyRs clustered at calcium release sites (CRSs). Close association of RyRs was observed in cardiac dyads (Asghari et al., 2014) as well as in purified cardiac RyRs (Cabra et al., 2016). However, only a small fraction of RyRs, whether in situ or in vitro, physically interact (Zahradnikova et al., 2010a; Jayasinghe et al., 2018). The first estimates of the junctional SR surface area per RyR were close to 900 nm^2^, i.e., 30×30 nm (Franzini-Armstrong et al., 1999; Soeller et al., 2007). The more recent estimates report values in the range of 3000 - 12500 nm^2^ (Baddeley et al., 2009; Hayashi et al., 2009; Hou et al., 2015; Galice et al., 2018; Jayasinghe et al., 2018). The average number of RyRs per CRS, often used as a measure of the CRS size, was estimated in the range of 8 – 22 for dyads at surface sarcolemma (Baddeley et al., 2009; Jayasinghe et al., 2018), and 8 – 100 for dyads at the tubular sarcolemma (Soeller et al., 2007; Hayashi et al., 2009; Scriven et al., 2010; Hou et al., 2015; Galice et al., 2018). Existence of RyR clusters separated by the edge-to-edge gap of less than 100 - 150 nm led to postulation of “superclusters” (Baddeley et al., 2009; Hou et al., 2015; Macquaide et al., 2015). Thus, the definition of CRS size is somewhat fuzzy. However, the question how the CRS size, the RyR number, and the RyR density affect calcium sparks is of fundamental physiological importance, since its resolving may help to understand variability of dyadic structure and function during postnatal development (Ziman et al., 2010; Louch et al., 2015; Mackova et al., 2017) and in various pathologies (Song et al., 2006; Wu et al., 2012). Understanding of dyads can be advanced by mathematical modeling, which allows exploration of the whole range of experimental data through the parametric space of defined models.

In this modeling study, we addressed the issues of calcium spark variability related to geometrical factors of RyR arrangement at calcium release sites. We developed a set of models allowing comparison of calcium release events produced by CRSs with different RyR number, arrangement, surface density, and single-channel calcium current amplitude. The model parameters spanned the whole range of relevant findings published recently. Activation of ryanodine receptor channels was simulated by a gating model conforming to the experimentally observed sensitivity of RyR2 to activation by Ca^2+^ ions and to inhibition by Mg^2+^ ions (Laver et al., 1997; Zahradnikova et al., 2003). We explain, on the grounds of the mass-action law, how synergy of CRS variables affects properties of local calcium release events.

## Methods

Simulations: Programs for generating the topology of CRSs and for simulating the activity of RyRs at calcium release sites were written in C++ and compiled in g++ as previously described (Iaparov et al., 2019). Both programs were parallelized using OpenMP. Parallel generation of random numbers was performed using OMPRNG (Bognar, 2013).

Computations were performed on the computer cluster URAN (Ural Branch of the Academy of Sciences of the Russian Federation).

Analyses: Data analysis was performed in OriginPRO Ver. 2019b (OriginLab, USA). Equations were fitted by the Levenberg-Marquardt method. Significance of differences between distributions was calculated using the Kolmogorov-Smirnov test.

### Model formulation

Each calcium release site, CRS, was represented by a set of RyRs characterized by geometrical and functional descriptors. The geometrical descriptors included number of RyRs, surface area per RyR, CRS density type, arrangement patterns of RyR distribution, and coordinates of RyRs. The functional description included amplitudes of RyR single-channel calcium current, and rate constants of the RyR gating model. The CRS models included calcium buffers for description of calcium diffusion (Iaparov et al., 2019). These descriptors of CRS, except for the rate constants and calcium buffering, were varied in the range of published values, and their impact on generation of calcium release events was evaluated.

#### The model of RyR gating

Simple two-state RyR gating models were used in previous studies to minimize requirements for computational power (Groff and Smith, 2008; Walker et al., 2014; Walker et al., 2015). These models generate the probability of the closed-to-open state transitions proportionally to concentration of the free Ca^2+^ ion ([Ca^2+^]) at the RyR center, according to the Hill model of mass action (Figure 1). These models inherently assume zero open probability at zero calcium concentration. Moreover, they underestimate RyR open probability at low Ca^2+^ concentration (Tencerova et al., 2012) and overestimate RyR open probability at high [Ca^2+^] for neglecting the effect of Mg^2+^ binding to the RyR inhibition sites (Laver et al., 1997; Zahradnikova et al., 2003). To overcome these problems, we used a 2 state model with open probability dependent on free Ca^2+^ and Mg^2+^ concentration(Iaparov et al., 2019). The open state probability *P*_*O*_ for the reversible closed (C) to open (O) state transitions in the two-state model:

**Figure 1.**
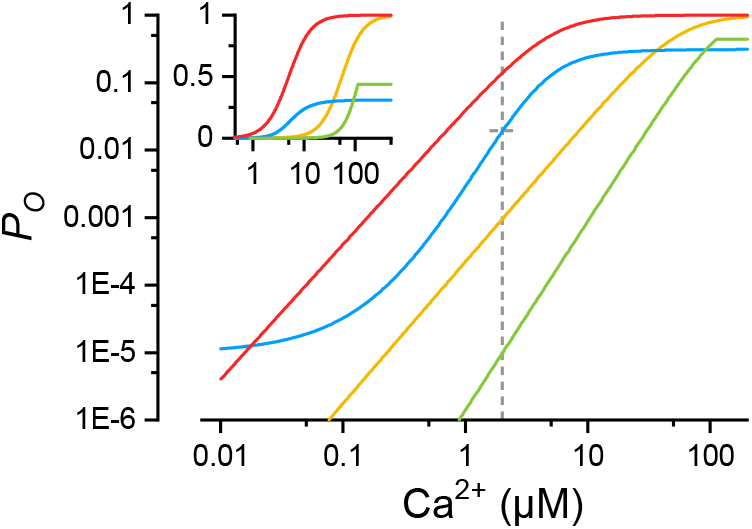
Calcium dependence of RyR open probability for two-state gating models. Blue line – this study, calculated by Eqs. 2 and 3. Red line - Groff and Smith (2008). Green line - Cannell et al. (2013). Yellow line - Walker et al. (2015). The dashed line indicates the iso-concentration level used in Figure 2. The inset shows the same data with the ordinate in linear scale.

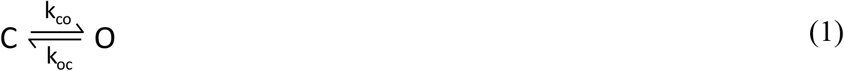

was described by:

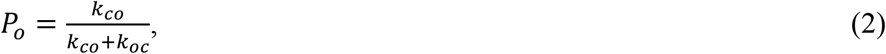

where *k*_*oc*_ = 1/15 ms^-1^ is the off-rate constant corresponding to the mean RyR open time of 15 ms estimated at a realistic luminal Ca^2+^ concentration in the absence of Cs^+^ ions (Gaburjakova and Gaburjakova, 2006), and *k*_*co*_ is the on-rate constant, dependent on [Ca^2+^] and [Mg^2+^] as:

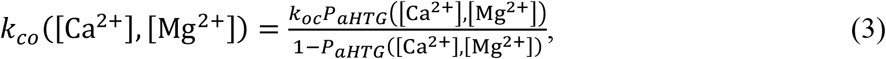

where *P*_*aHTG*_([Ca^2+^], [Mg^2+^]) is the steady-state open probability of the full aHTG model (Zahradnikova et al., 2010b; Iaparov et al., 2019). The [Ca^*2*+^] and [Mg^2+^] are the free ion concentrations calculated for the center of the RyR. The resulting *P*_*O*_ dependence on [Ca^2+^] is given in Figure 1 (blue line).

#### Generation of calcium release sites

Four topologically different arrangement patterns of calcium release sites were made by positioning 10, 20, 40, or 80 RyRs on the CRS plane (Figure 2), to encompass the range of reported CRS sizes and RyR surface densities (Baddeley et al., 2009; Hou et al., 2015; Galice et al., 2018; Jayasinghe et al., 2018). The number of RyRs (*N*_*RyR*_) and the CRS area per RyR (*A*_*C/R*_) defined the size and the density type of resulting CRS. The side *a* of square CRSs was calculated as 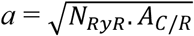.

**Figure 2.**
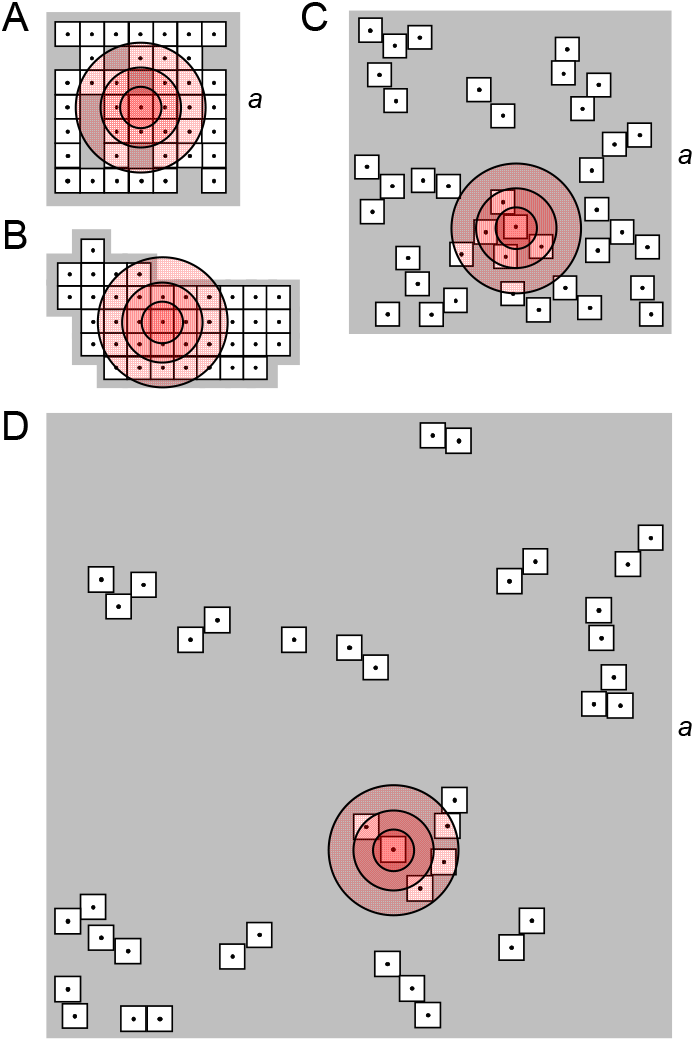
Geometric models of calcium release sites of different density types. The CRSs were selected from sets with *N*_*RyR*_ = 40 generated for the given CRS density type. **A:** A compact CRS with RyRs arranged side-by-side on a 7×7 lattice with *a* = 210 nm. **B:** A typical compact CRS with RyRs arranged side-by-side according to Walker et al. (2015). **C:** A narrow CRS with RyRs distributed according to pattern B with *a* = 374 nm. **D:** The wide CRS with RyRs distributed according to pattern B (Table 1) with *a* = 707 nm. The shadowed areas represent the planes of CRSs. The empty squares with dots represent RyRs with a central ion channel, the source of Ca^2+^ flux. The concentric circles of increasing diameter depict 2 μM [Ca^2+^] levels calculated for *i*_*Ca*_ of 0.04, 0.15, and 0.4 pA, at which the relative open probability of a RyR would be 2% (see Figure 1).

The geometric characteristics of CRSs were specified for the three CRS density types as follows:

**Table 1.** Pattern composition of narrow and wide calcium release sites.

| Pattern | CRS composition |  |  |  |
| --- | --- | --- | --- | --- |
|  | Chb | SbS | Both | Isl |
| **A** | 0.821 | 0.051 | 0.013 | 0.115 |
| **B** | 0.500 | 0.400 | 0.067 | 0.033 |
| **C** | 0.165 | 0.691 | 0.041 | 0.103 |

Chb, SbS, Both, Isl - checkerboard, side-by-side, both Chb and SbS, and isolated RyRs pattern, respectively. The data are average fractions that correspond to Asghari et al. (2014). The exact number of Chb, SbS, Both and Isl channels for CRSs of different size is given in Iaparov et al. (2019)

#### *<u>The compact type CRSs</u>* were of two variants

The square variants of compact CRSs (Figure 2A) contained the given number of RyRs placed randomly on a rectangular grid of 30×30 nm squares. The resulting *A*_*C/R*_ was 1440, 1125, 1103, and 911 nm^2^ per RyR, respectively. A set of five compact square CRSs was generated for each *N*_*RyR*_ (altogether 20 CRSs). The irregular variants of compact CRS (Figure 2B) used the experimentally determined coordinates (Walker et al., 2015) of 9 - 96 RyRs placed on a grid with 31-nm spacing. Altogether 107 of these CRSs had *A*_*C/R*_ equal to 961 nm^2^ per RyR.

#### The narrow type CRSs

(Figure 2C) had RyR density corresponding to that estimated by Jayasinghe et al. (2018). All narrow CRSs had *A*_*C/R*_ of 3481 nm^2^ (equivalent to a square of 59×59 nm per RyR). For each N_RyR_, the set of 100 CRSs of different RyR coordinates was generated (altogether 400 narrow CRSs).

#### The wide type CRSs

(Figure 2D) had RyR density corresponding to that estimated by Galice et al. (2018). All wide CRSs had *A*_*C/R*_ of 12544 nm^2^ (equivalent to a square of 112×112 nm per RyR). For each N_RyR_, the set of 100 CRSs of different RyR coordinates was generated (altogether 400 wide CRSs). Arrangement of RyRs on the CRS plane in the compact CRSs followed the side-by-side pattern (Yin et al., 2005; Groff and Smith, 2008; Walker et al., 2015). The topology of the narrow and the wide CRSs consisted of RyRs arranged into three patterns with defined proportions of neighbors in the checkerboard configuration, in the side-by-side configuration, and in isolation, as observed by electron microscopy (Asghari et al., 2014) under different experimental conditions (Table 1). Pattern A was dominated by RyRs in the checkerboard configuration, typical for low Mg^2+^ concentrations. Pattern C was dominated by RyRs in the side-by-side configuration, typical for high Mg^2+^ concentrations. The balanced pattern (B) contained the checkerboard, the side-by-side, and the isolated RyR configurations, as observed for near physiological conditions. The placement of RyRs in the narrow and wide CRSs according to patterns A, B or C was solved as an optimization problem (Iaparov et al., 2017; Iaparov et al., 2019) by genetic simulated annealing (GSA; (Xu et al., 2011)). The resulting RyR coordinates were used to calculate the set of inter-RyR distances.

#### Simulation and analysis of CREs

The variable distribution of RyRs among dyads observed experimentally was reflected by generating a large set of geometrical models of CRSs that complied with experimental data on clustering of RyRs into distinct patterns and on the number and density of RyRs in dyads. Each CRS construct was subjected to simulations of calcium sparks, in which the stochastically simulated activity of all RyRs in response to the initial opening of one of RyRs was monitored for a time interval of 200 ms. The stochastic nature of RyR gating gave rise to temporal fluctuations in the number of open RyRs. Therefore, each simulation of the CRS activity, initiated by a single random opening of one of RyRs, was repeated 10000 times for each CRS construct. Thus, each RyR of the CRS was initialized by 10000/*N*_*RyR*_ times on average.

The amplitude of calcium current through a single RyR, *i*_*Ca*_, or through a single CRS, were not yet determined *in situ*. Numerous experimental difficulties make the estimates ambiguous because of their dependence on many variables that are difficult to control in experiments. Janicek et al (2012) concluded that the calcium release flux per single RyR should be near 0.4 pA, the value satisfying estimates by independent experiments. In previous modelling studies of Ca sparks, much lower estimates were used; 0.04 pA (Groff and Smith, 2008), or 0.15 pA (Walker et al., 2015). To unveil the impact of single RyR calcium current amplitude on spark generation in CRSs of different type, we performed simulations using four *i*_*Ca*_ amplitudes of 0.04, 0.15, 0.25, and 0.4 pA. The *i*_*Ca*_ amplitude stayed constant during the simulation.

Concentration of calcium ions inside the CRS is not homogenous (Valent et al., 2007; Walker et al., 2014). At each locality, it depends on positions of open RyRs and on the single channel calcium current in a complex manner, being affected by diffusion rates, geometry of the CRS, buffering by mobile and immobile buffers, and calcium binding kinetics. In the CRS simulations, calcium concentrations were calculated for each RyR position, based on the given *i*_*Ca*_ value, on the distance from the source RyR, and on steady-state linearized buffered calcium diffusion (Naraghi and Neher, 1997; Iaparov et al., 2019), following each opening or closing of any RyR in the CRS. A record of the number of open RyRs in time *N*_O_(*t*) was generated using the Monte Carlo approach and the Gillespie algorithm.

The simulated traces of RyR activity were characterized by the maximal number of simultaneously open RyRs (*N*_*O*_ ^*peak*^), by the time to peak (*TTP*), and by the relative peak amplitude, *A*_*rel*_, calculated as:

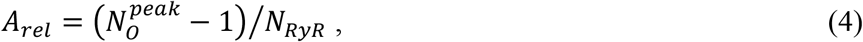

where (*N*_*O*_ ^*peak*^ – 1) is the number of RyRs in a CRS recruited at the time of the peak. These parameters were used for construction of histograms and further analyses as described below. Kinetic parameters of calcium release events were estimated from averaged traces.

### Vicinity and state of calcium release sites

According to the calcium induced calcium release theory, Ca^2+^ ions are the interaction element between RyRs. This fact was handled in different ways in published models of CRSs. For CRSs with uniform RyR arrangement, Williams et al. (2011) introduced the mean-field approximation of RyR-RyR interaction. This approximation is not applicable to a general case of non-uniform arrangement of RyRs (Iaparov et al., 2019). For compactly packed RyRs on a regular lattice, Walker {, 2014 #78} used the maximum eigenvalue of the RyR adjacency matrix as an overall parameter characterizing RyR arrangement. The adjacency principle, however, cannot be used to characterize freely distributed RyRs. Other measures of RyR arrangement, namely the average nearest-neighbor distance, the four-nearest-neighbor distances, and the mean RyR occupancy, all related to the density and the mean RyR-RyR distance in CRSs (Cosi et al., 2019), correlated with parameters of simulated calcium sparks relatively weakly. Recently, we have found that the maximum eigenvalue of the RyR inverse distance matrix is a promising predictor of spark fidelity for general CRSs {Iaparov, 2017 #2}; however, it is somewhat abstract and difficult to perceive. Therefore, we define here a new geometric parameter that characterizes the spatial disposition of RyRs in a general CRS more conceivably, and as demonstrated in the Results, allowing to interpret simulated calcium release events on common grounds.

The steady-state solution of the diffusion equation relates the Ca^2+^ distribution to the positions of open RyRs. In an unbuffered system, the concentration of Ca^2+^ ions at individual RyR positions is proportional to the sum of reciprocal values of distances *D*_*ij*_ to the open RyRs, 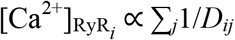 (Crank, 1975). Note that the presence of calcium buffers decreases the steady-state concentration of Ca^2+^ ions in the CRS (Naraghi and Neher, 1997), thus the above relationship between RyR geometry and [Ca^2+^] is an upper bound for the RyR-RyR interaction through Ca^2+^ ions. The inverse distance can be thought of as the closeness *C*_*ij*_ = 1/*D*_*ij*_, where indices *i* and *j* correspond to elements of the RyR distance matrix. This means that formation of a calcium release event by interacting RyRs should be related to the RyR closeness matrix **C** composed of elements *C*_*ij*_, where the elements of *i*^*th*^ row correspond to closeness of the remaining RyRs (*j* ≠ *i*) to the *i*^*th*^ RyR. Let’s consider the process of increasing the area of a square in such a way that all coordinates will be multiplied by a factor *K*. This process represents a linear transformation on a surface that transforms the distances between two points on the surface in proportion to the factor *K*, and the surface area in proportion to the square of *K*. Thus, when the surface area of a CRS, given as *S*_*CRS*_ = *A*_*C/R*_ × *N*_*RyR*_, is increased by a factor *K′*, the distances between RyRs are increased proportionally to the square root of *K′*, i.e., 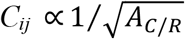. We propose that an analogous relationship can be formulated when *S*_*CRS*_ is changed by changing the number of RyRs and keeping *A*_*C/R*_ constant, i.e., 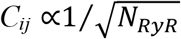 Considering these relationships allows to define a new parameter that does not change with *N*_*RyR*_ - the RyR vicinity *v* of the *i*^*th*^ RyR to the *j*^*th*^ RyR - as the normalized closeness 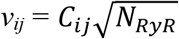. The effect of Ca^2+^ flux through the *i*^*th*^ RyR on the whole CRS can then be considered a function of the average vicinity *v*_*i*_ of the *i*^*th*^ RyR to the remaining RyRs (*N*_*RyR*_ -1) defined as:

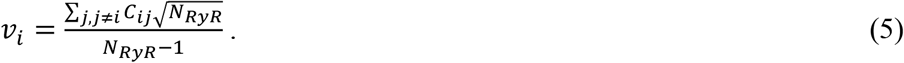

The whole calcium release site can be then characterized by the average vicinity of all its RyRs, i.e., the group RyR vicinity *v*:

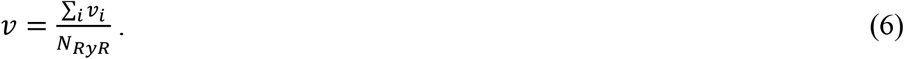

The group RyR vicinity defined in this way thus represents the CRS state variable characterizing the spatial component of the RyR-RyR interaction potential. The usefulness of this characterization will become apparent later.

Since CRS activity upon opening of the *i*^*th*^ RyR depends also on the amplitude of calcium current through this RyR, we can define a state variable *φ*_*i*_ pertinent to and characterizing the *i*^*th*^ RyR as the weighed product of calcium current and RyR vicinity:

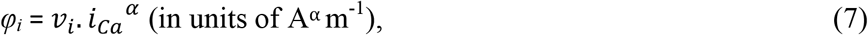

where the power *α* is the weighing factor for relative contributions of calcium current and vicinity of the *i*^*th*^ initiating RyR to the CRS response. Developing this idea to account for the start of CRS activation by opening of any of its RyRs, the group RyR vicinity *v* (Eq. 6) allows definition of the group RyR state variable, i.e., the CRS state variable *φ* as:

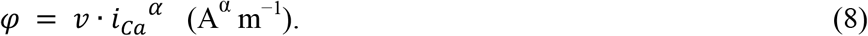

The values of the exponent *α* were found by relating *φ*_*i*_ or *φ* to the characteristics of simulated calcium release events by means of state-response functions:

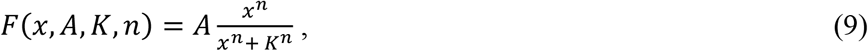

where *x* stays for *φ*_*i*_ or *φ, A* is the amplitude, *K* is the half-maximum value of *x*, and *n* is the slope factor.

## Results

This study surveys calcium release events generated by 10108 CRS models (2527 CRSs at four *i*_*Ca*_ values) covering the range of published RyR densities, sizes and arrangements, as described in Model formulation. A CRS was defined by the set of its properties: the number of RyRs (*N*_*RyR*_ of 9 - 96), the density type (compact, narrow, or wide; Figure 2), the arrangement of RyRs into specific patterns in the case of narrow and wide CRSs (A, B, or C; Table 1), and the single-channel calcium current amplitude *i*_*Ca*_. The compact, narrow and wide CRSs were generated with density of 1 RyR per 911-1440, 3500, and 12500 nm^2^, respectively. The effect of the single-channel RyR calcium current amplitude was inspected at 0.04, 0.15, 0.25, and 0.40 pA. CRE simulation was initiated by opening of one of RyRs in the CRS, and recorded for 200 ms. The CRE records were characterized by their peak amplitudes, expressed as the maximal number of simultaneously open RyRs, their relative amplitude, and their time to peak. The CRS activity was characterized by the average relative amplitude and by the fraction and average time course of different response types (see below) calculated from 10000 records.

### Basic characteristics of simulated calcium release events

Calcium release sites responded to the initial RyR opening by calcium release events of three types of (Figure 3):

**Figure 3.**
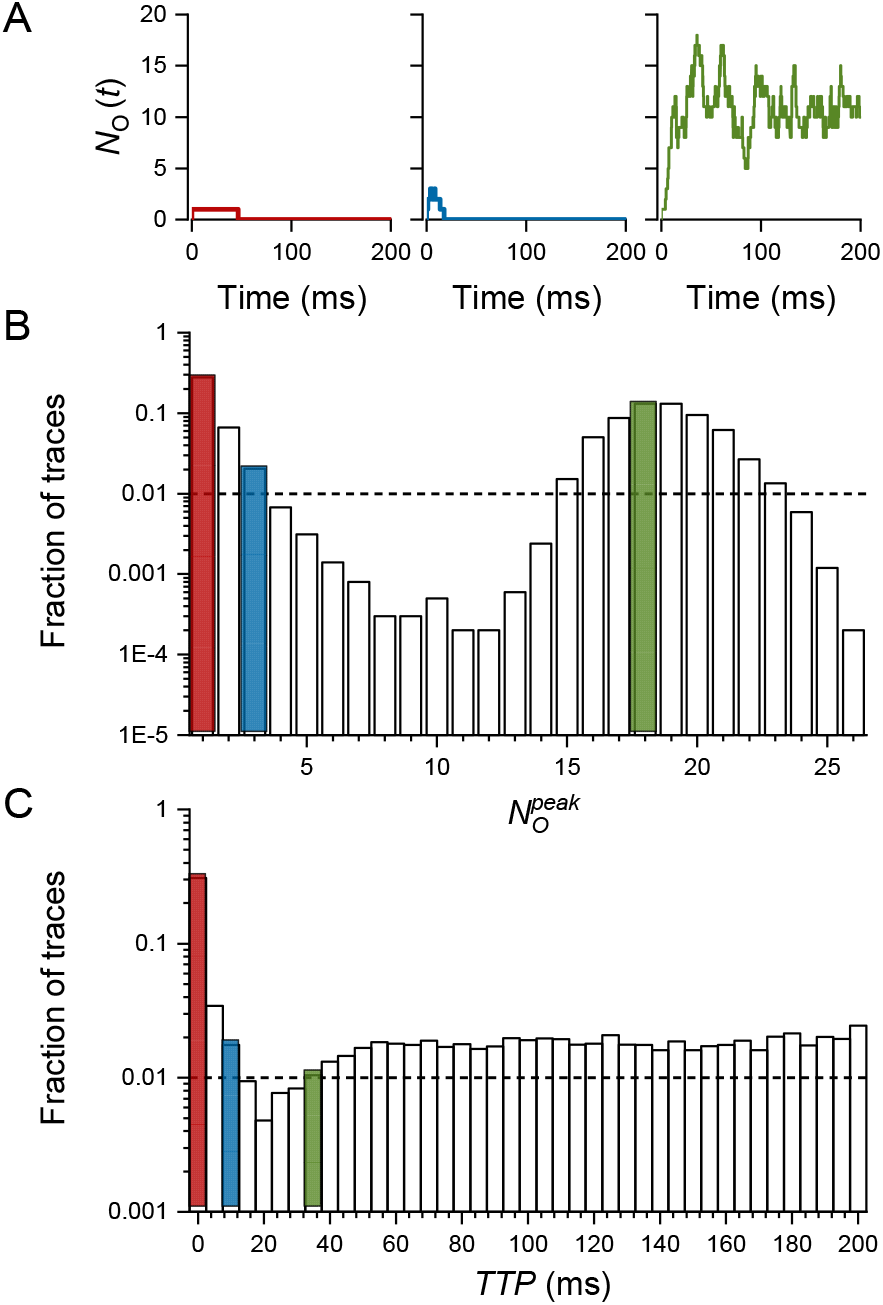
Types and characteristics of calcium release events. The exemplified dataset consisted of 10000 CREs, generated in a narrow CRS of pattern B, 40 RyRs, and *i*_*Ca*_ of 0.4 pA. The color-coded histogram bins correspond to traces of the same color. **A**: Typical traces of three types of CREs: a quark (left), a blip (center), and a spark (right). Arrowheads indicate *N*_*O*_^*peak*^ and time to peak (*TTP*) positions. **B:** The distribution of peak amplitudes (*N*_*O*_^*peak*^) in the dataset. Quarks, blips and sparks represented 27.9 %, 10.0 %, and 62.1 % of traces, respectively. The nadir of the histogram was at 11 simultaneously open RyRs. **C**: The distribution of *TTP* values in the dataset. The dashed lines in B and C denote the occurrence level of 1 %; note the logarithmic scales on the ordinates.

- The responses that consisted solely of the initial RyR opening, which did not trigger other RyR openings, were classified as quarks (Lipp and Niggli, 1996; Brochet et al., 2011; Wescott et al., 2016). Peak amplitudes of quarks were equal to one open RyR, and durations followed exponential distribution with a time constant somewhat less than the RyR mean open time.
- The responses containing openings of multiple RyRs, which peaked rapidly and then spontaneously deactivated, were classified as blips (not related to the type of IP3R calcium transients (Swillens et al., 1998)). Peak amplitude of blips was small, typically in the interval between 2 and the nadir value of the amplitude histogram. Blips typically peaked in less than 10 ms and ceased in less than 100 ms. It should be noted that the amplitude of individual blips could reach up to 50% of *N*_*RyR*_ in compact CRSs; however, since individual blips peaked at different times, their average peak amplitude was only 1.2 - 1.6 open RyRs.
- The responses containing openings of many RyRs, with time to peak typically of >20 ms, and with RyR openings occurring until the end of the 200-ms simulation, were classified as sparks. Their peak amplitude was larger than the nadir of the amplitude histogram, and their time to half amplitude was typically about 15 ms.

The fraction of quarks, blips, and sparks (*F*_*q*_, *F*_*b*_, and *F*_*s*_, respectively) were determined as the amplitude of the first bin (*N*_*O*_^*peak*^ = 1), the sum of bin amplitudes from 2 to the nadir, and the sum of bin amplitudes at and above the nadir, respectively.

The amplitude histogram in F gure 3B shows pronounced bimodal distribution where 99 % of records contained either less than 4 simultaneously open RyRs about 37%) or more than 14 simultaneously open RyRs (about 62%). The time-to-peak histogram (Figure 3C), also with bimodal distribution, indicates that the fraction of short events (about 30%) corresponded to the fraction of quarks and blips (Figure 3B), and that about 70% of short e**v**ents peaked in less than 10 ms. It also indicates that a majority of events peaked between 35 - 200 ms with almost uniform distribution, as expected for random fluctuations in the number of simultaneously open RyRs.

### Amplitude distribution of calcium release events depends on CRS determinants

The distribution of peak ampl tudes of simulated calcium release events varied substantially with the number of RyRs, with the CRS **d**ensity type, and with he single-channel calcium current amplitude (Figure 4). The quarks and blips prevailed when the RyR number, density and *i*_*Ca*_ were low. Spark type events dominated when at least one of these parameters was high. When *i*_*Ca*_ was low (0.04 pA, Figure 4, red bars), the sparks activated rarely and compact CRSs with the largest *N*_*RyR*_ (Figure 4, bottom left).

**Figure 4.**
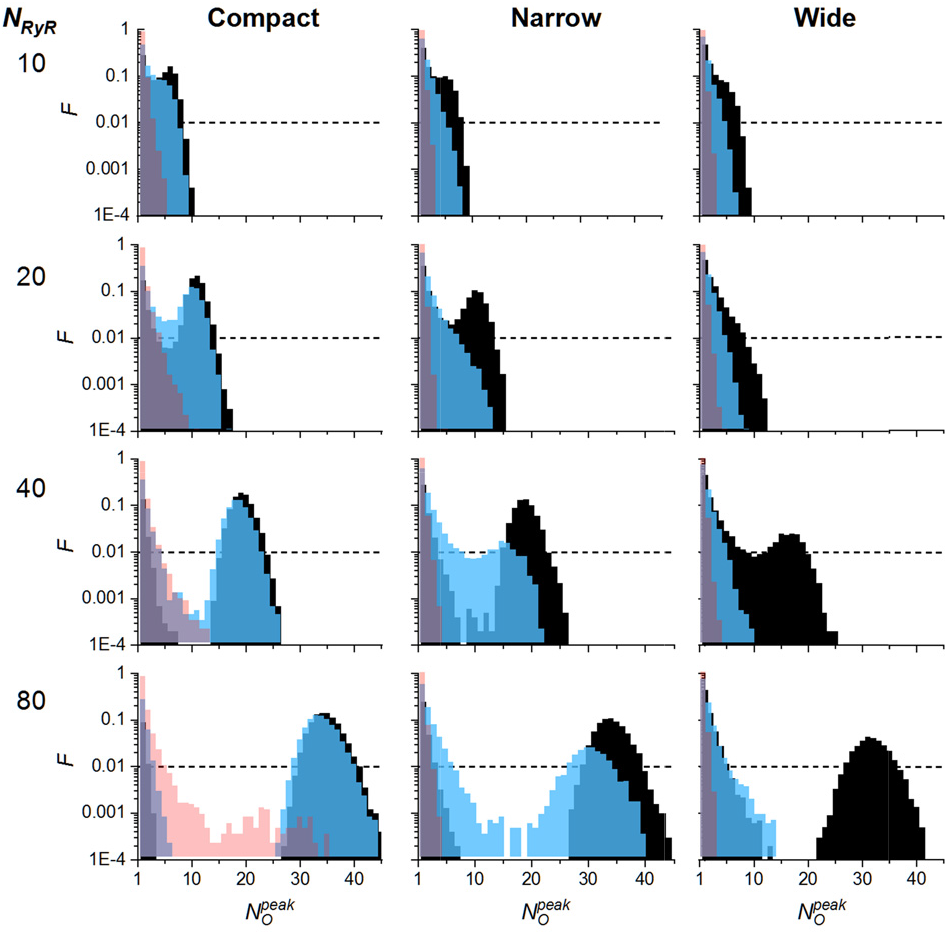
Typical distributions of peak amplitudes of calcium release events. Calcium release sites of various density type, *N*_*RyR*_, and *i*_*Ca*_ are co mpared. The black, blue and red bins correspond to CREs generated by *i*_*Ca*_ of 0.40, 0.15 and 0.04 pA, respectively. The dashed lines denote occurrence level of 1 %. The exemplified datasets consisted of 10000 simulated CREs generated for one typical CRS of each type a nd size. The narrow and wide CRS types were of pattern B. *F* – fraction of traces. Note the logarithmic scale on the ordinates.

For a majority of simulated conditions, the amplitude histograms showed bimodal distribution reflecting the large difference in the amplitudes of blips and sparks. The bimodal behavior indicates the existence of a critical or thr**e**shold condition, attributable to the extent of calcium increase during the rising pha e of the ca**l**cium release event. If the effective [Ca^2+^] increase was subthreshold under the given conditions, it might have failed to activa**t**e more RyRs, since the released calcium quickly dissipated and the release event terminated due to stochastic extinction. In small CRSs, the amplitude distribution was not clearly bimodal, so that the distinction between blips and sparks was not so prominent. The peak number of simultaneously open RyRs in individual **s**parks did not reach *N*_*yR*_ but it highly exceeded the limit expected for the given steady-state open probability of the gating mo**d**el (*P*_*O*_ = 0.31).

These *N*_*O*_^*peak*^ excursions were due to re-openings of RyRs. On average, about 4 re-openings per RyR occurred in the 200-ms simulations. The maximal number of simultaneously open RyRs reached the full number of RyRs in the CRS (*N*_*RyR*_) only in small CRSs (10 RyRs). In large CRSs (80 RyRs), the *N*_*O*_^*peak*^ reached only about 55 % of RyRs even at the largest *i*_Ca_ (0.4 pA) and in the densest CRSs. At lower *i*_*Ca*_ values or larger inter-RyR distances, *N*_*O*_^*peak*^ values were much lower than the number of RyRs in the given CRS. This was due to the steep [Ca^2+^] gradient in the buffered milieu of the CRSs and low [Ca^2+^] at peripheral RyRs, resulting in long closed times of RyRs distant from the open RyRs.

### CRS activity depends on distribution of RyRs

The strength of Ca^2+^ mediated interaction between RyRs depends on distances between RyRs, which may have different distribution in different CRSs of the same RyR density. The distribution of RyRs in CRS could be conveniently characterized by means of their group vicinity (Eq. 6), which increases with the extent of RyRs clustering. The overall vicinities calculated for the whole range of generated CRSs are given in Table 2. At the same number of RyRs, the group RyR vicinity of various CRS types differed to the extent that their ranges almost did not overlap. The group vicinity increased with *N*_*RyR*_ in compact CRSs, but did not change in narrow and in wide CRSs. This was due to different *S*_*RyR*_ in the compact CRSs but constant *S*_*RyR*_ in the other two CRS types. Nevertheless, the group vicinities for individual CRS of the same type and of the same *N*_*RyR*_ also varied, as evidenced by their ranges, but now due to the randomness of RyR spatial distribution. Thus, the group RyR vicinity distinguishes well between CRSs with different RyR spatial distribution.

**Table 2.**
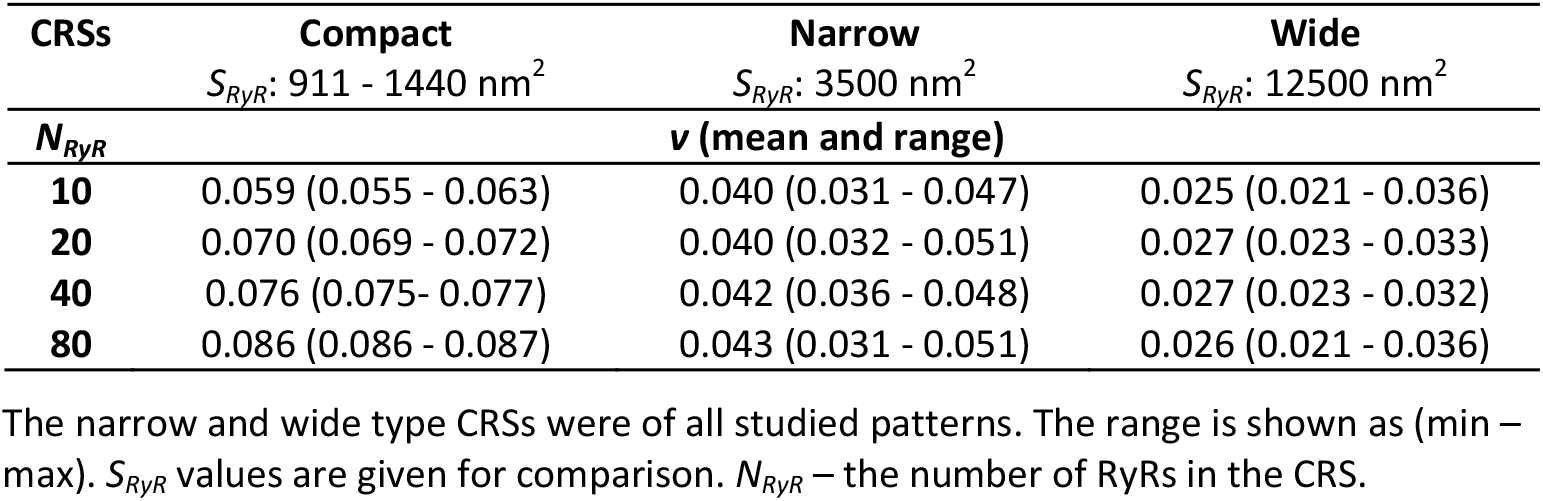
Overall RyR vicinities in CRS models of different density type (Eq. 2).

| CRSs | Compact<br>$S_{RyR}$: 911 - 1440 nm<sup>2</sup> | Narrow<br>$S_{RyR}$: 3500 nm<sup>2</sup> | Wide<br>$S_{RyR}$: 12500 nm<sup>2</sup> |
| --- | --- | --- | --- |
| $N_{RyR}$ | $\nu$ (mean and range) | | |
| 10 | 0.059 (0.055 - 0.063) | 0.040 (0.031 - 0.047) | 0.025 (0.021 - 0.036) |
| 20 | 0.070 (0.069 - 0.072) | 0.040 (0.032 - 0.051) | 0.027 (0.023 - 0.033) |
| 40 | 0.076 (0.075 - 0.077) | 0.042 (0.036 - 0.048) | 0.027 (0.023 - 0.032) |
| 80 | 0.086 (0.086 - 0.087) | 0.043 (0.031 - 0.051) | 0.026 (0.021 - 0.036) |

The narrow and wide type CRSs were of all studied patterns. The range is shown as (min – max). $S_{RyR}$ values are given for comparison. $N_{RyR}$ – the number of RyRs in the CRS.

Comparison of simulated responses of individual CRSs of identical density and pattern type but of different RyR distribution (Figure 5) revealed that the calcium release events provided different amplitude distributions. The difference between the two histograms in Figure 5, showing distribution of event amplitudes for two narrow CRSs of pattern B, was highly significant (P = 1.6 × 10^−76^). The more “clustered” CRS (Figure 5, blue) generated 23 % of quarks, 7 % of blips and 70 % of sparks. The less “clustered” CRS, with group RyR vicinity smaller by 12 %, was substantially less active (Figure 5, red). It generated more quarks (31 %) and blips (11 %) but less sparks (58 %). Thus, the arrangement of RyRs in the CRS has an influence on calcium release activity *via* the extent of clustering between RyRs, and the group RyR vicinity could be considered a good integral descriptor of RyR clustering.

**Figure 5.**
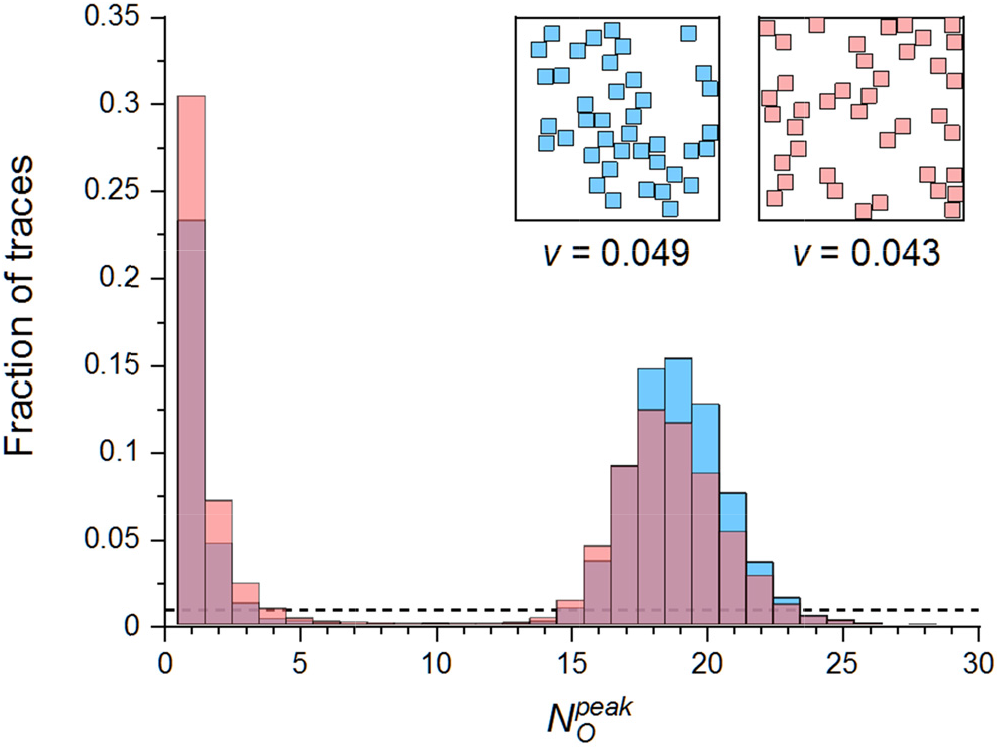
Comparison of amplitude distribution of calcium release events of two calcium release sites differing in RyR vicinity. The exemplified datasets consisted of 10000 simulated CREs per CRS at *i*_*Ca*_ of 0.40 pA. Two narrow type CRSs of pattern B and 40 RyRs, shown in the insets, represen ed the most (blue) and the least (red) clustered RyRs in the set of 300 narrow CRS; Kolmogorov-Smirnov test for difference between distributions: P = 1.6 × 10^−76^. The dashed line denotes o**c**currence level of 1%. Note the linear scale on the ordinate. The quarks contributed to bin 1, the blips to bins 2 - 10, and the sparks **c**ontributed to bins ≥11 of *N*_*O*_ ^*peak*^ values.

### CRS activity does not depend on RyR pattern type

We tested whether arrangement of RyRs into checkerboard and side-by-side patterns in the experimentally observed proportions A, B, C (Table 1) would generate physiologically meaningful differences in CRS activity. For narrow CRSs and *i*_*Ca*_ of 0.40 pA, the relative occurrences of quarks, blips, and sparks showed only minor dif erences between different patterns with the same *N*_*RyR*_ (Table 3). These differences were much smaller than those between CRSs of the same pattern type but different *N*_*RyR*_. Vari tion of the RyR patterns in wide CRSs, and/or at smaller *i*_*Ca*_ amplitudes, did not produce substantial differences in CRS activity either (data not shown). Note that RyRs of the compact CRSs are always arranged in the side-by-side pattern. Nevertheless, for the given pattern type, the fraction of sparks increased with increasing *N*_*RyR*_, while the fractions of quarks and blips showed an opposite trend (Table 3). The values of group RyR vicinity were the same for all *N*_*RyR*_ values.

**Table 3.** Dependence of fractions of event types on RyR patterns.

| | $N_{RyR}$ | Pattern A | Pattern B | Pattern C |
| --- | --- | --- | --- | --- |
| $F_q$ | 10 | $41.6 \pm 0.3$ | $42.1 \pm 0.3$ | $45.2 \pm 0.3$ |
| | 20 | $32.8 \pm 0.2$ | $33.3 \pm 0.2$ | $34.9 \pm 0.2$ |
| | 40 | $27.5 \pm 0.1$ | $27.3 \pm 0.1$ | $28.2 \pm 0.1$ |
| | 80 | $24.1 \pm 0.1$ | $23.9 \pm 0.1$ | $24.3 \pm 0.1$ |
| $F_b$ | 10 | $25.1 \pm 0.1$ | $25.7 \pm 0.2$ | $26.7 \pm 0.1$ |
| | 20 | $17.9 \pm 0.2$ | $18.8 \pm 0.2$ | $20.1 \pm 0.2$ |
| | 40 | $9.6 \pm 0.1$ | $9.7 \pm 0.1$ | $10.3 \pm 0.1$ |
| | 80 | $6.5 \pm 0.0$ | $6.3 \pm 0.0$ | $6.5 \pm 0.0$ |
| $F_s$ | 10 | $33.4 \pm 0.4$ | $32.2 \pm 0.5$ | $28.1 \pm 0.4$ |
| | 20 | $49.3 \pm 0.3$ | $47.9 \pm 0.3$ | $45.0 \pm 0.3$ |
| | 40 | $62.9 \pm 0.2$ | $63.0 \pm 0.2$ | $61.4 \pm 0.2$ |
| | 80 | $69.4 \pm 0.1$ | $69.8 \pm 0.1$ | $69.2 \pm 0.1$ |

Narrow CRSs; $i_{Ca} = 0.40$ pA. All values are given as mean ± SD. $F_q$ – fraction of quarks; $F_b$ – fraction of blips; $F_s$ – fraction of sparks. $F_q + F_b + F_s = 1$ .

These results do not indicate importance of the arrangement pattern *per se*. This could be expected, since the only factor controlling activation of RyRs in these CRS models was the local calcium concentration. However, the absence of an effect of side-by-side or checkerboard arrangement patterns on CRS activity does not mean that biological factors affecting arrangement of RyRs into patterns do not affect CRS behavior. It only means that the pattern alone plays a negligible role, since it affects only the nearest RyR to RyR distances. In real dyads, however, a change in the pattern could indicate a change in the degree of RyR clustering, that is, a change in the RyR vicinity. In fact, this is very realistic situation. Let consider a hypothetical experiment, in which some biological factor, e.g., phosphorylation or Mg^2+^, causes aggregation of RyRs in dyads (Asghari et al., 2014). Then, presumably, the re-arrangement of RyRs from the mostly random to the mostly side-by-side or mostly checkerboard distribution will generate clusters with increased RyR vicinity. The dyad could be morphologically the same but its CRS density type would change. For instance, a single large wide CRS will change to two narrow or even to four compact smaller CRSs. The changes in activity calculated for this scenario are given in Table 4. These dramatic changes in the dyad behavior underline the impact of RyR arrangement on frequency and amplitude of spontaneous sparks.

**Table 4.** Effect of dyad composition on relative occurrence of event types.

|  | One wide CRS | Two narrow CRSs | Four compact CRSs |
| --- | --- | --- | --- |
| Spark type | 80 RyRs | 2 x 40 RyRs | 4 x 20 RyRs |
| Quarks | 0.44 | 0.29 | 0.17 |
| Blips | 0.27 | 0.10 | 0.07 |
| Sparks | 0.29 | 0.61 | 0.76 |

$i_{Ca} = 0.40$ pA. The three dyads, each 80 RyRs, were made of 1, 2, or 4 CRSs by changing the pattern of RyR clustering. The dyads were activated by opening of 1 randomly selected RyR in one of CRSs at a time. The activation of one CRS did not affect other CRSs in the dyad. Wide and narrow CRSs were of pattern B.

### CRS activity relates quantitatively to CRS state variables

The relationship of CRS activity to CRS variables, i.e., to RyR surface area, RyR number, and *i*_*Ca*_ amplitude, are summarized in Figure 6 for narrow and wide CRSs. These results extend the indicative findings obtained for a single CRS per condition (see above). The fraction of quarks decreased with increased *N*_*RyR*_; the decrease was more prominent for narrow CRSs than for wide CRSs (Figure 6, red segments). As counterpart, the fraction of sparks increased with increasing *N*_*RyR*_ (Figure 6, blue segments). Note that the sparks did not occur at the lowest *i*_*Ca*_ in any of the CRSs. In wide CRSs, the increase in the fraction of sparks with *N*_*RyR*_ was apparent only at the highest *i*_*Ca*_ values. The fraction of blips was substantial in all CRSs. In wide CRSs it dominated over sparks at any *N*_*RyR*_ and *i*_*Ca*_. The fraction of blips varied with the CRS determinants in a complex manner with specific features for each set of conditions (Figure 6, green segments). The decrease of *i*_*Ca*_ amplitude led to a prominent decrease in the fraction of sparks, and complementarily, to an increase in the fraction of blips and especially of quarks, for any *N*_*RyR*_ in both CRS density types. The substantial difference in the behavior of narrow and wide CRSs can be attributed to the limited outreach and high steepness of calcium gradient within a CRS (see Figure 2) leading to higher recruitment of more clustered RyRs (see below).

**Figure 6.**
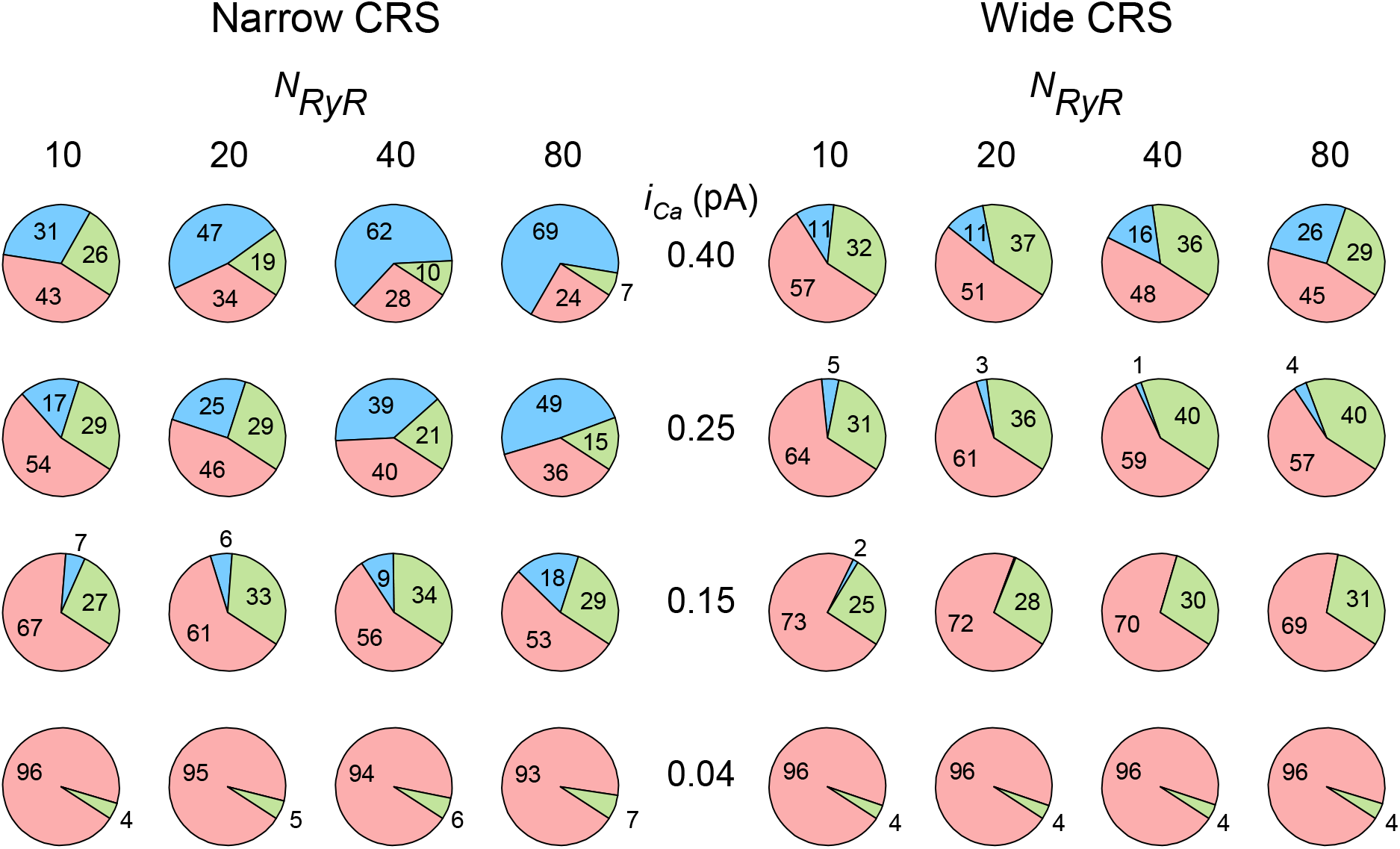
Relative occurrences of different calcium release event types. Summary for different *N*_*RyR*_ and *i*_*Ca*_ in narrow and wide calcium release sites. Fractions of sparks, blips and quarks are shown as blue, green, and red segments, respectively, and indicated by numbers (in %). Data represent averages of 10000 simulated records per CRS, calculated for all 300 geometric CRS models under each condition.

The above analysis revealed how the CRS activity is controlled by variables defining the CRS state (Figures 3 and 5). However, the parameters of calcium release events did not have a clear dependence on either the area per RyR, or the number of RyRs per CRS, or the single-channel amplitude of calcium current. There was a clear dependence of *Ā* _*rel*_ on the group RyR vicinity; albeit the dependence was different for different *i*_*Ca*_ values (Figure 7A). In contrast, all characteristics of calcium release events correlated well with the CRS state variable *φ* (Figure 7B). The values of *φ* varied between 0.002 and 0.05 pA^0.67^ nm^−1^ in the whole simulation dataset across all wide, narrow, compact, and contact-network CRSs. The fractional occurrence of quarks, *F*_*q*_, declined to 0.1 for *φ* values around 0.045 pA^0.67^ nm^−1^. The fractional occurrence of blips, *F*_*b*_, showed a maximum of about 0.4 at small *φ* values between 0.01 and 0.02, and was negligible at *φ* above 0.04 pA^0.67^ nm^−1^. The fractional occurrence of sparks, *F*_*s*_, began to dominate at *φ* above 0.02 pA^0.67^nm^−1^. The relationship between the average relative amplitude *Ā*_*rel*_ and *φ* showed half-maximum at *φ* of about 0.02 and maximum of about 0.4 at *φ* above 0.04 pA^0.67^ nm^−1^.

**Figure 7.**
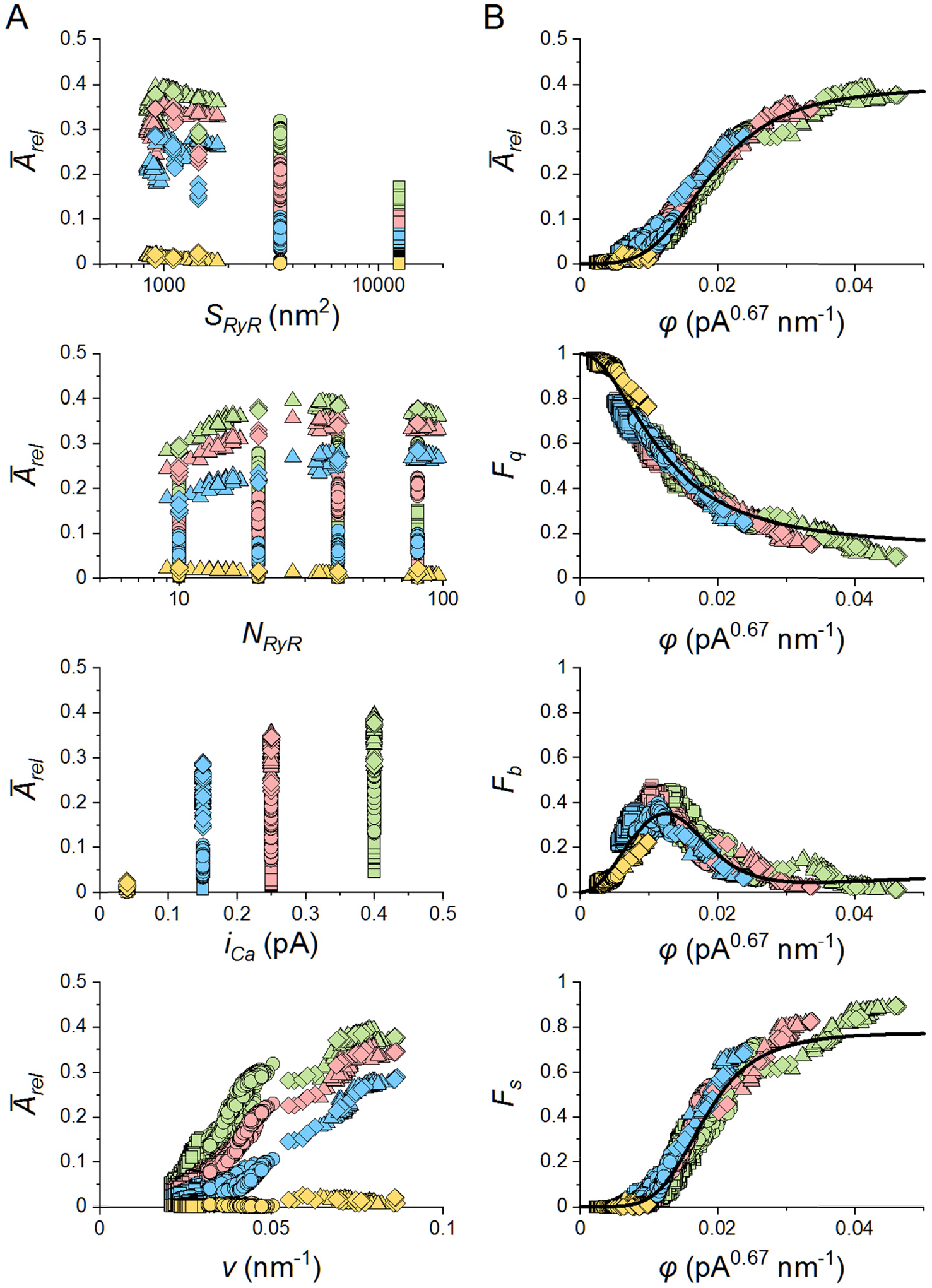
Relationships between characteristics of calcium release events and of calcium release sites. **A -** Relationships between average relative event amplitudes *Ā*_*rel*_ a nd determinants of CRSs. *S*_*RyR*_ – the area per RyR; *N*_*RyR*_ – the number of RyRs per CRS; *i*_*Ca*_ – the single-channel calcium current. *ν* – the group RyR vicinity. **B -** Relationships between characteristics of calcium release events and the CRS state variable *φ. F*_*q*_ – fracti on of quarks. *F*_*b*_ – fraction of blips. *F*_*s*_ – fraction of sparks. *Ā*_*rel*_ - average relative event amplitud e. The lines show the respective best global fit of all data points by functions derived from E qs. 8 and 9 (*F*_*q*_ = 1 - *F*(*φ,A*_*1*_,*K*_*1*_,*n*_*1*_), *F*_*b*_ = *F*(*φ,A*_*1*_,*K*_*1*_,*n*_*1*_) - *F*(*φ,A*_*2*_,*K*_*2*_,*n*_*2*_), *F*_*s*_ = *F*(*φ,A*_*2*_,*K*_*2*_,*n*_*2*_), *Ā* _*rel*_ = *F*(*φ,A*_*3*_,*K*_*3*_,*n*_*3*_), the indices 1, 2, 3 correspond to indices *i* in Table 5 that lists the parameters of the f unctions. In A and B, yellow, blue, red, and green symbols denote data for *i*_*Ca*_ values of 0.0 4, 0.15, 0.25, and 0.40 pA, respectively. Squares and circles denote data for all wide and narro w CRSs (1200 geometric models per CRS density type containing 10 - 80 RyRs). Triangles and diamonds denote data for the two types of compact CRSs (20 and 107 geometric models, respectively, containing 9 - 96 RyRs).

The dependences of CRE characteristics on the CRS state variable *φ* were approximated by respective functions derived from Eq. 8 and 9. The best fit parameters summarized in Table 5 were found by relating all observed event characteristics of all simulated CRSs to the variable *φ* in parallel. The data characterizing CRSs of the compact type at large values of *i*_*Ca*_ showed small systematic deviations in *F*_*b*_ and *F*_*s*_ (green triangles and diamonds in Figure 7B). These deviations occurred in small CRSs (*N*_*RyR*_ ≤ 20), which contained a large fraction of RyRs at edges and corners, and for which the amplitude distributions did not show a clear nadir.

**Table 5.** The best fit parameters of the CRE dependences on the CRS state variable ϕ.

| $\alpha = 0.67 \pm 0.002$ | | | | |
| --- | --- | --- | --- | --- |
| | $i$ | $A_i$ | $K_i$ | $n_i$ |
| $1-F_q$ | 1 | $0.87 \pm 0.003$ | $0.012 \pm 5E-5$ | $2.07 \pm 0.01$ |
| $F_s$ | 2 | $0.78 \pm 0.002$ | $0.018 \pm 5E-5$ | $4.87 \pm 0.02$ |
| $\bar{A}_{rel}$ | 3 | $0.40 \pm 0.004$ | $0.019 \pm 1E-4$ | $3.69 \pm 0.05$ |

The functional relationship between the CRS state variable *φ* and the characteristics of calcium release events across their types and across the CRS density types for the whole range of *N*_*RyR*_ and *i*_*Ca*_ values (Figure 7B) indicate that the *φ* has qualities of a descriptor of the CRS state. The *φ* value predicts the source propensity of a CRS to produce sparks and blips upon spontaneous opening of a random RyR, in terms of their probability of occurrence and average amplitude. From the point of physical meaning, the variable *φ* could be considered as an effective calcium concentration in the dyadic gap, since it consolidates the characteristics of stochastic calcium release events (Eq. 9).

### Kinetics of release events is a function of the CRS state variable *φ*

Kinetics of calcium release events of different type varied substantially, as demonstrated in Figure 8. The termination rate of averaged quarks (Figure 8A, top row) was shorter than the closing rate of the RyR gating model, since it was curtailed by activation of blinks and sparks that increased in occurrence with the CRS variable *φ*. As a result, its decay time (Figure 8B, top row) was a decreasing Hill function of *φ* with a maximum of 15 ms equal to the mean RyR open time, and with *K* = 0.014 ± 2.9E-4, and *n* = 1.75 ± 0.06 estimated from 48 averaged records.

**Figure 8.**
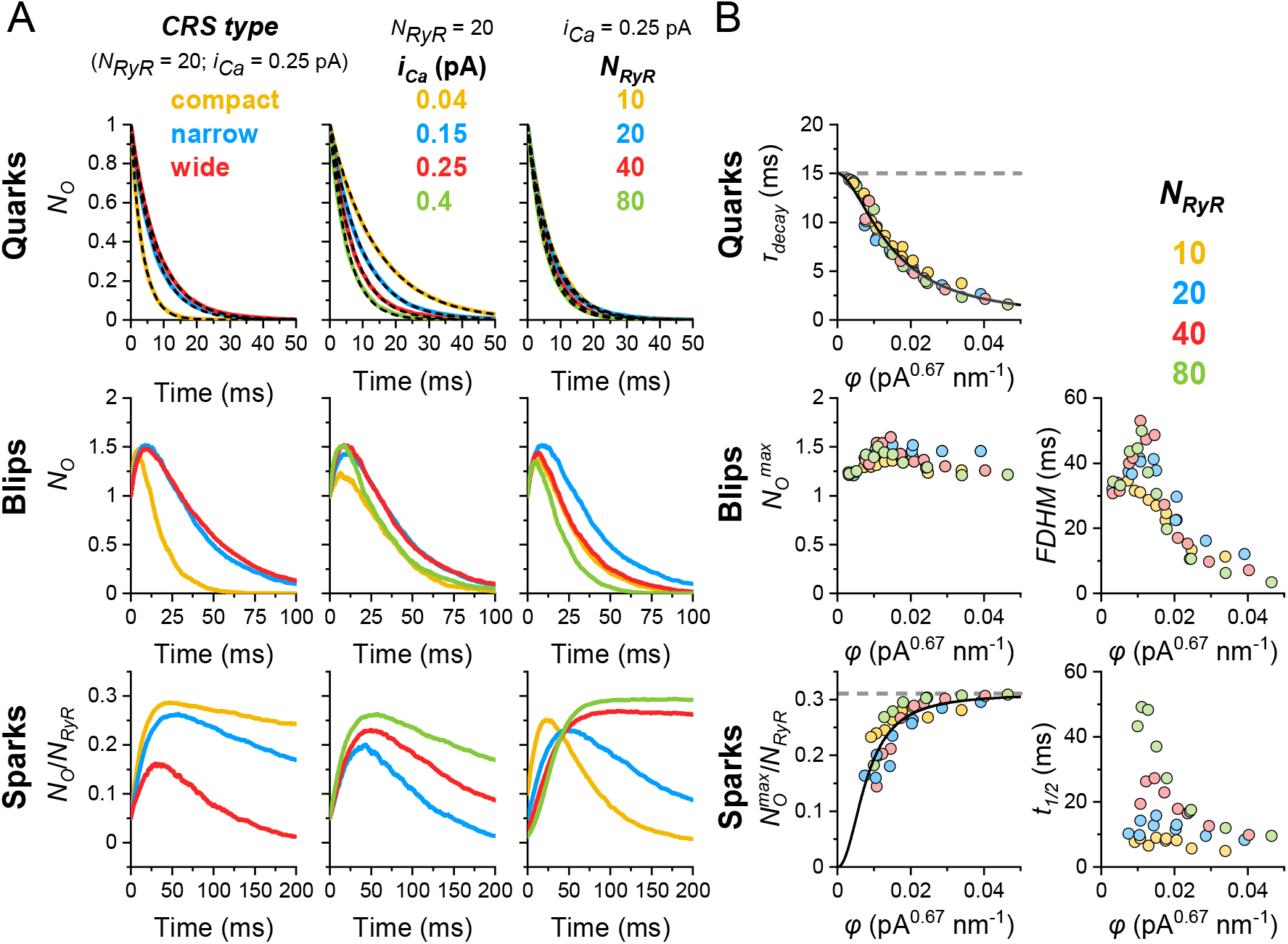
Kinetics of averaged calcium release events in calcium release site models. The data was obtained from 10000 simulated CREs generated for one typical CRS of each type and size. The narrow and wide CRS types were of pattern B. **A** - The average time courses of quarks, blips and sparks. Left: compact, narrow and wide CRSs with *N*_*RyR*_ = 40 and *i*_*Ca*_ = 0.4 pA. Center: narrow CRSs with *N*_*RyR*_ = 40 and of the indicated range of *i*_*Ca*_ values. Right: narrow CRSs with *i*_*Ca*_ of 0.40 pA and the indicated range of *N*_*RyR*_. The time course of quarks was fitted by single exponentials (dashed lines). **B** - Characteristics of the averaged CREs. Quarks: The time constants of quark decay were approximated by a decreasing Hill function of *φ* (*A*- *F*(*φ,A,K,n*), where *F*(*φ,A,K,n*) is defined by Eqs. 8 and 9, with *A* equal to the RyR open time, *K* = 0.014 ± 2.9E-4 and *n* = 1.75 ± 0.06). Blips: *N*_*O*_^*max*^ and the FDHM of the average time courses of blips. Sparks: *N*_*O*_^*max*^/*N*_*RyR*_ and the half-time of activation of averaged time courses of sparks. *N*_*O*_^*max*^/*N*_*RyR*_ was approximated by an increasing Hill function of *φ* (Eqs. 8 and 9, with *A* equal to the maximum *P*_*O*_ of RyR, *K* = 0.0079 ± 6.1E-4 and *n* = 2.13 ± 0.30).

The time course of averaged blips was always biphasic (Figure 8A, middle row). Kinetic parameters were dependent on the CRS variable *φ* (Figure 8B, middle row). The values of *TTP* and *FDHM* increased in the *φ* range of 0 - 0.01 pA^0.67^nm^−1^ due to the increased recruitment of RyRs and then declined at higher *φ* due to faster formation of sparks (see below). The maximal values of kinetic parameters were 12 and 53 for *TTP* and *FDHM*, respectively.

The time course of average sparks (Figure 8A, bottom row) depended on the CRS variable *φ*, but their amplitude was proportional to *N*_*RyR*_. As a result, the maximum of the ensemble open probability was an increasing Hill function (Eq. 9) of *φ* with parameter *A* = 0.31 equal to the maximum RyR open probability, and with K = 0.0079 ± 6.1E-4, and n = 2.13 ± 0.30. Average sparks of CRSs with *N*_*RyR*_ ≤ 20 had a clearly defined peak with TTP < 70 ms in the whole

range of *φ*. In contrast, average sparks of CRSs with *N*_*RyR*_ ≥ 40 had clearly defined TTP only for *φ* < 0.15 pA^0.67^nm^−1^ (data not shown); at higher *φ* values their ensemble open probability stayed constant until the end of simulation. For CRSs with *N*_*RyR*_ ≥ 40, the value of activation halftime, t_*1/2*_, reached a peak around *φ* = 0.01 pA^0.67^nm^−1^ and then declined again, indicating faster conversion of blips into sparks. This is apparent from the similar dependence of the FDHM of blips and t_*1/2*_ of sparks on *φ* (Figure 8B).

### Effectiveness of spark initiation depends on the initiating RyR

In simulation studies on contact network CRSs, Walker et al. (2014; 2015) showed that the efficiency of spark activation depends on the position of the initiating RyR. A substantial increase in spark fidelity was observed only if the initiating RyR channel was located in the core of the CRS, since RyRs at the periphery had less neighbor RyRs than RyRs in the core of the CRS. Although a similar phenomenon could be expected in less dense CRSs as well, the extent of RyR recruitment in CRSs of different internal disposition could not be predicted. Therefore, we analyzed recruitment of RyRs by an initiating RyR at all positions in the CRS for the defined range of CRS variables (*N*_*RyR*_, *S*_*RyR*_, and *i*_*Ca*_). The initiating RyR, characterized by its RyR state variable *φ*_*i*_, affected distinctly the relative occurrence of release events and the average relative amplitude of CRS responses, 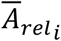, as summarized in Figure 9. It is apparent that differences between RyRs with higher and lower levels of local clustering, i.e., with higher and lower vicinity, become more apparent in CRSs with lower group RyR vicinity (insets in individual panels of Figure 9). A typical compact, narrow and wide CRS with *N*_*RyR*_ of 10, 20, 40 and 80 RyRs per CRS was analyzed for the whole range of studied *i*_*Ca*_ amplitudes (0.04, 0.15, 0.25, and 0.40 pA). Compact CRSs with 10 RyRs, which were the source of the outliers in Figure 7B, were not included in this analysis. Altogether 11 CRSs and 440 RyRs were examined. The dependences of calcium release event characteristics on *φ*_*I*_ are shown in Figure 9. The average relative amplitude 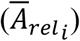 and fraction of sparks 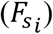 increased, while the fraction of quarks 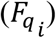 decreased with increasing *φ*_*i*_. The fractional occurrence of blips 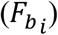 showed a maximum of about 0.4 at *φ*_*i*_ ≈ 0.01 pA^0.76^ nm^-1^. The dependences of calcium release event characteristics on *φ*_*i*_ were approximated by respective functions derived from Eqs. 8 and 9 as follows: Relative amplitude 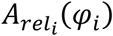 and the fraction of sparks 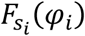 were described directly by *F(<p*_*i*_, *A, K, n)*, the fraction of quarks was described by 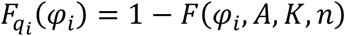, and the fraction of blips was described by 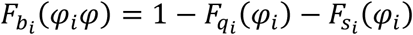. The value of the exponent *α* was found by relating all observed event characteristics of all simulated CRSs to the variable *φ*_*i*_ in parallel. The best fit parameters are summarized in Table 6.

**Table 6.**
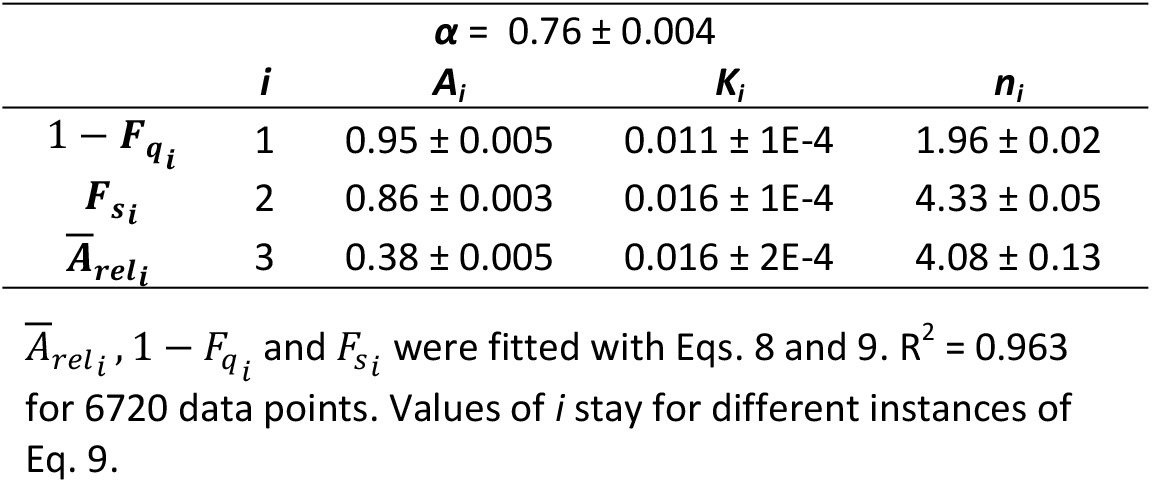
Parameters of CRE dependence on the RyR_*i*_ state variable *φ*_*i*_.

**Figure 9.**
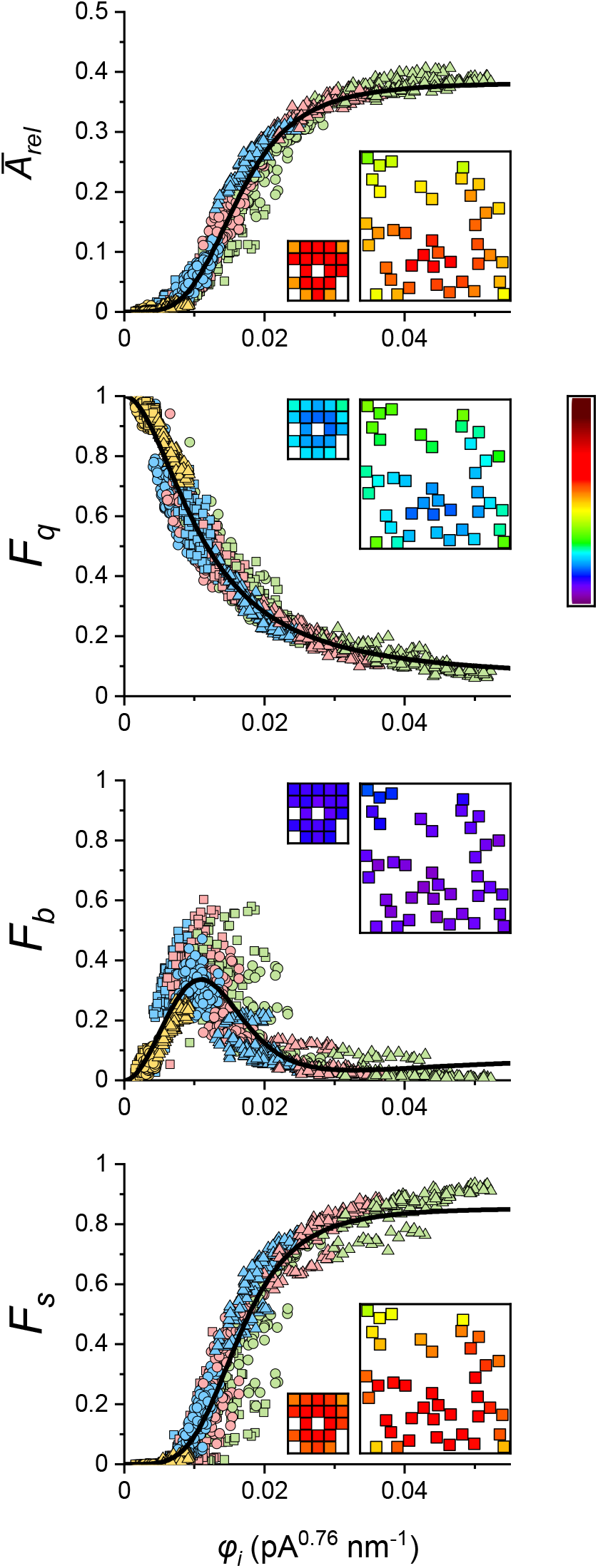
Relationships between characteristics of calcium release events and the RyR state variable *φ*_*i*_ of the initiating RyRs. *Ā*_*rel*_ - average relative CRE amplitude. *F*_*q*_ – fraction of quarks. *F*_*b*_ – fraction of blips. *F*_*s*_ – fraction of sparks. Yellow, blue, red, and green symbols denote results for *i*_*Ca*_ values of 0.04, 0.15, 0.25, and 0.40 pA, respectively. The squares, circles and triangles represent wide, narrow, and compact CRSs, respectively. The lines show the respective best global fit of all data points by functions derived from Eqs. 8 and 9 (*F*_*q*_ = 1 - *F*(*φ*_*i*_,*A*_*1*_,*K*_*1*_,*n*_*1*_), *F*_*b*_ = *F*(*φ*_*i*_,*A*_*1*_,*K*_*1*_,*n*_*1*_) - *F*(*φ*_*i*_,*A*_*2*_,*K*_*2*_,*n*_*2*_), *F*_*s*_ = *F*(*φ*_*i*_,*A*_*2*_,*K*_*2*_,*n*_*2*_), *Ā* _*rel*_ = *F*(*φ*_*i*_,*A*_*3*_,*K*_*3*_,*n*_*3*_)),where the indices 1, 2, 3 correspond to indices *i* in Table 6 that lists the parameters of the functions. Insets: the exemplar compact CRS with 20 RyRs at *i*_*Ca*_ = 0.25 pA and the exemplar narrow CRS with 40 RyRs at *i*_*Ca*_ = 0.4 pA, both showing (average) response values for *n* = 1000 simulations for individual initiating RyRs, color-coded according to the look-up table bar (right) scaled for respective ordinates.

The RyR state variable *φ*_*i*_ allowed to outline relationships valid across spark characteristics and CRS density types at different *N*_*RyR*_ and *i*_*Ca*_. From the point of physical meaning, the variable *φ*_*i*_ could be considered an agent representing the effective calcium concentration around the individual RyR in the dyadic gap.

### CRS spark activity results from synergy between RyR vicinity and *i*_*Ca*_

A prediction of the model is the existence of a functional relationship between spark activity and the CRS state variable *φ*, and between spark activity and the RyR state variable *φ*_*i*_. The existence of these state variables is the result of synergy between the calcium source strength component (*i*_*Ca*_^*α*^) and the topological component (group RyR vicinity) of the CRS environment. The common dependence of the relative occurrence of individual event types (quarks, blips and sparks), as well as of their relative amplitude on *i*_*Ca*_ and ν is shown in Figure 10. The amplification of the effect of *v* by *i*_*Ca*_ and vice versa is manifestation of the synergy between *i*_*Ca*_ and *v*.

**Figure 10.**
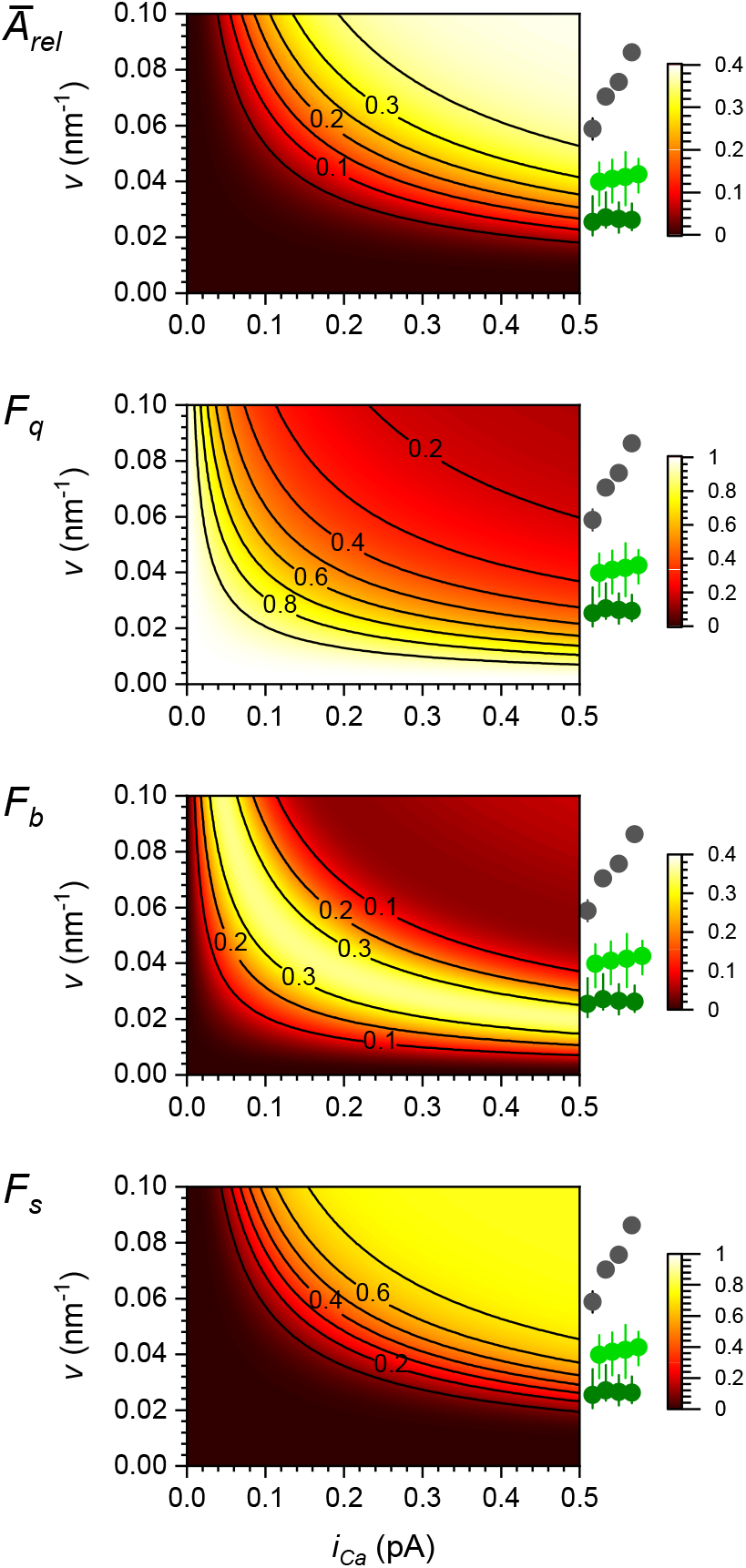
Synergy graphs between group RyR vicinity and single channel calcium current, for relative occurrences and amplitudes of calcium release events. The calculations were performed using Eqs. 8 and 9 with values from Table 5; the calculated values of the respective variables (symbols on the left) are color coded according to the look-up table bars (right side of the figure rows) and marked by iso-lines. The symbols to the right of each plot indicate means and ranges of group vicinities for CRSs models used in this study (black - compact CRSs; vivid green - narrow CRSs; dark green - wide CRSs) calculated for *N*_*RyR*_ of 10, 20, 40 and 80 (from left to right, respectively).

## Discussion

This study focused on the role of geometric determinants of cardiac dyads in formation of elementary calcium release events. We used a modelling approach that allowed wide-range exploration of dyad-defining factors implemented to numerous CRS models, which would not be feasible in real experiments. The scope was limited to a realistic range of factors, as determined up to now. To keep computational costs reasonable, a simple model of RyR gating that, however, provided realistic calcium dependence of RyR open probability was used together with a simple model of buffered calcium diffusion between RyRs. The *in silico* experiments, emulating spontaneous calcium sparks initiated by opening of a single RyR in CRS, generated random calcium release events that were recorded as a time sequence of RyR openings and analyzed. Under majority CRS conditions, three types of calcium release events, resembling elementary CREs, known as quarks, blips and sparks, were generated. Analysis of their relative occurrence, amplitude and kinetics revealed high cooperativity between the number, geometry, and single-channel calcium current of RyRs in formation of CRS responses. These results were explained by introduction of the CRS state variable defined as weighted contribution of microscopic CRS determinants - the single-channel calcium current and the vicinity of RyRs in CRS, where the vicinity is a function of the number of RyRs in the CRS and of their mutual distances.

Recent experimental and theoretical studies established the importance of dyad structure and RyR clustering on spontaneous calcium spark formation (Cannell et al., 2013; Walker et al., 2014; Walker et al., 2015; Kolstad et al., 2018; Cosi et al., 2019) as well as on triggered calcium release (Zahradnik et al., 2013; Cosi et al., 2019; Novotova et al., 2020). In this study we used a large set of CRS geometries that approximated the structural data by constrained random distribution of RyRs in CRS (Iaparov et al., 2017; Iaparov et al., 2019). The CRS geometries followed the known concept of RyR arrangement into checkerboard, side-by-side, and isolated random patterns (Asghari et al., 2014), a realistic range of RyR number between 8 - 96, and CRSs with a range of RyR surface densities between of 900 - 12 500 nm^2^ per RyR (Baddeley et al., 2009; Hayashi et al., 2009; Hou et al., 2015; Galice et al., 2018; Jayasinghe et al., 2018). This imparted the known variability of dyadic RyR distribution to the inspected CRS models.

All geometric CRS models responded to a random RyR opening by calcium release events of amplitudes and time courses that were highly variable. The variability was high in CRSs of any geometry, despite the single-channel current amplitude being constant during calcium release events, and thus could be attributed to stochastic gating and group behavior of RyRs. Characteristics of CREs were, however, substantially modulated by the geometric disposition of RyRs. CREs of all three types were generated by each CRS model, although at different relative occurrences. Moreover, repeated CRS activation by the same initiating RyR, that is, at constant RyR vicinity, also produced all three types of CREs (Figures 4, 6). Therefore, we suppose that the various types of local calcium release events observed in cardiac myocytes under otherwise the same conditions represent a rudimentary variability of CRS responses. In other words, if experiments indicate broad range of local calcium release amplitudes (Wang et al., 2004; Zima et al., 2008; Brochet et al., 2011; Janicek et al., 2012) and durations, (Zima et al., 2008; Brochet et al., 2011) or even opening of one or a few RyRs (Wang et al., 2004; Janicek et al., 2012), it has roots in the theory of CRS operation. This modeling confirmed the predictions that calcium release may terminate even in the absence of a specific termination mechanism due to stochastic attrition (Stern et al., 1999). But it also predicts that a specific mechanism of calcium release termination should exist, since otherwise the typical duration of CRE would be much longer than physiologically meaningful.

We defined the CRS state variable *φ* as an internal determinant of CRS productivity, which combines the vicinity of interacting elements (RyRs) with the distribution of interaction agents (Ca^2+^ ions). It turned out that the stochastic, amplitude and kinetic characteristics of all three types of release events correlated well with the CRS state variable, but importantly, with the same value of the weight factor *α* for *i*_*Ca*_ (Figures 7B and 8). Therefore, the state variable *φ* could be thought of as the effective calcium concentration in the dyadic gap (Eq. 9), although its dimension A^*α*^.m^-1^ indicates linear current density, or calcium flux per distance or perimeter. Importantly, the same quantitative relationship satisfied responses of CRSs based on the experimentally determined (Walker et al., 2015) as well as on the algorithm-generated RyR distributions (Iaparov et al., 2019). Taken together, these multi-parametric correlations on the grounds of generalized RyR geometries support intuitive understanding that the vicinity and *i*_*Ca*_ of RyRs determine the activity of CRS in synergy (Figure 10).

This study introduced the concept of RyR vicinity (Eqs. 4 and 5), which characterizes distances between RyRs and the size of the RyR group by one numerical value. Unlike the RyR surface density or the number of RyRs in the CRS, the vicinity resolves the differences in distribution of RyRs in dyads. In other words, although two CRSs have the same number of RyRs and the same surface density, they have different CRS vicinity values. Likewise, two RyRs in the same or in different CRS will have different RyR vicinity value. To generalize, the vicinity is a measure of the RyR interaction potential since it increases with increasing RyR clustering. The previously used concept of adjacency or nearest neighborhood (Walker et al., 2014) attributes the interaction potential only to close neighbors, resulting in discretized levels of RyR-RyR interactions. In contrast to the RyR surface density or mean RyR occupancy used in previous models (Cannell et al., 2013; Cosi et al., 2019), the vicinity characterizes the whole CRS (Eq. 6), as well as individual RyRs (Eq. 5), according to their global or local geometric dispositions.

The state variables *φ* and *φ*_*i*_ based on vicinities *v* and *v*_*i*_, which deliver the synergic character of RyR clustering and of calcium current, allow to predict the relative occurrence of quarks, blips and sparks, as well as their amplitudes and kinetics in the whole range of studied CRS characteristics. However the predicton is in the stochastic sense, that is, valid on average. Individual responses of a CRS may vary broadly, as varies the chance of RyR opening. Taken from the other side, the CRS and RyR state variables determine the propensity of CRSs to generate specific type of calcium release events, their amplitude, and the time course. They may predict whether the specific CRS will generate preferentially quarks, blips, or sparks (Figure 10). In fact, the state variables make possible to explain the deterministic relation of stochastic processes of RyR gating and distribution, and of calcium diffusion and binding, to generation of calcium signals. To fully accomplish these goals, new models of CRS are needed that would better define the control of RyR gating by luminal calcium at both luminal (Terentyev et al., 2002; Laver, 2007; Tencerova et al., 2012) and cytosolic sites (Laver, 2007) and the RyR to RyR interaction by direct coupling (Stern et al., 1999; Marx et al., 2001).

### Physiological implications

The presented CRS models suggest that the decrease of *i*_*Ca*_ due to depletion of calcium in the junctional cisterna during calcium release may contribute to the termination of calcium sparks even in the absence of direct RyR regulation by luminal calcium (Terentyev et al., 2002; Laver, 2007; Tencerova et al., 2012). According to these results, spark activation is substantially slowed down and spark decay is substantially speeded up at decreased values of *i*_*Ca*_ (Figure 8). It is generally accepted that during a calcium spark a substantial local depletion of luminal Ca^2+^ concentration occurs (Brochet et al., 2005), amounting to ≈40% (Zima et al., 2008). In an example of a narrow CRS with 40 channels, a similar decrease in *i*_*Ca*_ would result in a reduction of the relative occurrence of sparks by 56% (from 0.55 to 0.24) at original *i*_*Ca*_ of 0.4 pA, or by 73% (from 0.26 to 0.07) at original *i*_*Ca*_ of 0.25 pA. Similar extrapolating considerations propose that dyadic SR cisternae with low Ca content or with lower RyRs Ca conductance may have a much lower probability of CRE activation.

Another mechanism of regulation of CRS activity is a change in RyR vicinity, which has a substantial effect on the fraction of sparks. For example, splitting of one wide 80-RyR CRS (*v* = 0.025) into two narrow 40-RyR CRSs (*v* = 0.04), then down to four compact 20-RyR CRSs (*v* = 0.06), and finally to eight compact 10-RyR CRSs (*v* = 0.08) would change fraction of sparks from 0.15 to 0.55, 0.73 and 0.76 at *i*_*Ca*_ = 0.40 pA, but from <0.01 to 0.07, 0.32 and 0.57 at *i*_*Ca*_ = 0.15 pA. The peak number of recruited channels in the four respective scenarios would be 14, 14, 9 and 4.6 per spark/blink at *i*_*Ca*_ = 0.40 pA, and 3.4, 4.7, 5.1 and 3.7 per spark/blink at *i*_*Ca*_ = 0.15 pA. Additionally, the decrease in vicinity results in slowing down of spark activation, in agreement with previous simulations and with experimental data (Kolstad et al., 2018).

### Limitations

To keep computational costs reasonable, a simple model of RyR gating that provided realistic calcium dependence of RyR open probability was used together with a simple model of buffered calcium diffusion between RyRs. Although it accounts for the inhibition by Mg^2+^ ions, this inhibition is carried out solely by decreasing the activation rate, which is not realistic (Zahradnikova et al., 2003).

This model of CRSs was built without a mechanism of calcium release termination, since the termination mechanism of calcium sparks would increase the complexity of CRS behavior that might blur the effect of geometrical factors, which were of prime interest in this study. For the same reasons we did not implement either coupled RyR gating mechanism or variability of single-channel calcium current in the course of CRE.

The use of the simplified model of buffered calcium diffusion in semi-infinite space (Naraghi and Neher, 1997) instead of calculating the full reaction-diffusion problem explicitly, and including the presence of the sarcolemmal membrane (Valent et al., 2007), substantially simplified and speeded up calculations but also reduced the buildup of calcium concentrations at individual RyRs. Although this simplification affected calculations of CRS activity quantitatively, we assume that its effects on the quality of correlations between CRE characteristics and the state variables or on the synergy between *i*_*ca*_ and vicinity were small, since activation of RyRs requires a much longer time than that necessary to reach a steady-state calcium gradient after opening a RyR in a CRS (Soeller and Cannell, 1997; Valent et al., 2007).

## Acknowledgments

The study was performed using the Uran supercomputer of the Krasovskii Institute of Mathematics and Mechanics. The research was supported by Russian Foundation for Basic Research (RFBR) according to the research project no 18-31-00153 (development of the computational model of calcium release site), by the Government of the Russian Federation, Program 02.A03.21.0006, by the Russian Ministry of Education and Science, project no FEUZ-2020-0054, by the Slovak Research and Development Agency project APVV-15-0302, by SAV-TUBITAK project JRP/2019/836/RyRinHeart, by the Science Grant Agency grant VEGA 2/0143/17, and by the Research & Development Operational Program funded by the ERDF (BIOMED, grant number ITMS: 26240220087).

## CRediT authorship contribution statement

Bogdan Iaparov: Conceptualization, Methodology, Software, Data Curation, Validation, Formal analysis, Investigation, Writing - original draft, Writing - review & editing, Project administration, Funding acquisition.

Ivan Zahradník: Conceptualization, Methodology, Validation, Investigation, Writing - original draft, Writing - review & editing. Visualization,

Alexander S. Moskvin: Conceptualization, Supervision, Writing - review & editing

Alexandra Zahradníková: Conceptualization, Methodology, Software, Data Curation, Validation, Formal analysis, Investigation, Writing - original draft, Writing - review & editing, Visualization, Supervision, Project administration, Funding acquisition.

